# Neural processing of communication signals: The extent of sender-receiver matching varies across species of *Apteronotus*

**DOI:** 10.1101/387324

**Authors:** K.M. Allen, G. Marsat

## Abstract

As communication signal properties change, through genetic drift or selective pressure, the sensory systems that receive these signals must also adapt to maintain sensitivity and adaptability in an array of contexts. Shedding light on this process helps us understand how sensory codes are tailored to specific tasks. In a species of weakly electric fish, *Apteronotus albifrons*, we examined the unique neurophysiological properties that support the encoding of electrosensory communication signals that the animal encounters in social exchanges. We compare our findings to known coding properties of the closely related species, *Apteronotus leptorhynchus*, to establish how these animals differ in their ability to encode their distinctive communication signals. While there are many similarities between these two species, we found notable differences leading to relatively poor coding of the details of chirp structure occurring on high-frequency background beats. As a result, small differences in chirp properties are poorly resolved by the nervous system. We performed behavioral tests to relate *A. albifrons* chirp coding strategies to its use of chirps during social encounters. Our results suggest that *A. albifrons* do not exchange frequent chirps in non-breeding condition, particularly when the beat frequency is high. These findings parallel the mediocre chirp coding accuracy in that they both point to a reduced reliance on frequent and rich exchange of information through chirps during these social interactions. Therefore, our study suggests that neural coding strategies in the central nervous system vary across species in a way that parallels the behavioral use of the sensory signals.

**SIGNIFICANCE:** Sender-receiver matching is a phenomenon commonly observed in the peripheral nervous system. It enables communication production and reception to evolve together so that conspecifics remain sensitive to important signals. In this manuscript we examine this phenomenon in the central nervous system of the weakly electric fish *A. albifrons* and compare its processing of communication signals to a closely related species (*A. leptorhynchus).* Although some differences across the two species can help tailor the system for processing species-specific signals, our data indicate that encoding of communication signals in *A. albifrons* is not as sensitive as in *A. leptorhynchus* for certain categories of signals. Our data support the idea that the extent of sender-receiver matching can vary as a function of behavioral needs.

## INTRODUCTION

Sender and receiver matching facilitating communication has been demonstrated across diverse groups, including songbirds (Brumm and Slabbekoorn, 2005), anurans (Schul and Bush, 2002), and insects (Neuhofer et al., 2008). Peripheral tuning and call matching are well characterized, and studies have shown evidence for sender and receiver matching in response to auditory coding of courtship song (Gerhardt and Schwartz, 2001; Tootoonian et al., 2012; Woolley and Moore, 2011) but little work has been done in other sensory modalities. We aim to identify species-specific variations in coding properties of central electrosensory neurons and link them to the divergent use of communication signals, showing that the extent of species-specific adaptations results in a variably tight sender-receiver match.

Weakly electric fish are ideal for examining diversification in signal production and reception. Apteronotids share a common mode of communication, modification of their electric organ discharge (EOD), but exhibit huge variety in signal structure. Voluntary modulations of EOD (chirps) vary dramatically in properties such as duration, frequency, and shape, even between closely related species (Turner et al., 2007; Zakon and Smith, 2002). Likewise, chirp reception is influenced by EOD waveform shape (Petzold et al., 2016), chirp features or structure (Benda et al., 2006; Marsat et al., 2012), and social environment (Stamper et al., 2010). The apteronotid electrosensory system could have coding properties generic enough to process these signals efficiently despite differences in structure and use between species; alternatively, differences in chirp processing could reflect adaptations of the electrosensory system to these differences in chirp production.

Relative EOD frequencies of interacting fish greatly influences chirp perception. These fish perceive ongoing amplitude modulations (AM beat) that are the product of two fish with different EOD frequencies interacting at close range (Bastian, 1981). Chirping modulates this beat (Hagedorn and Heiligenberg, 1985; Zupanc and Maler, 1993). In *A. leptorhynchus*, low frequency beats are typical of agnostic encounters between fish of similar sex and size, and fish respond with frequent small chirps (<100Hz increase) (Hupé and Lewis, 2008). Higher frequencies occur between fish of opposite sexes or with large differences in body size and more often elicit big chirps (>100Hz), typical of courtship (Hagedorn and Heiligenberg, 1985; Henninger et al., 2018).

Chirp coding has been well characterized in *A. leptorhynchus.* The two categories of signals described above produce different responses in the primary sensory area, the electrosensory lateral line lobe (ELL). Small chirps on low frequency beats cause stereotyped bursting among ELL pyramidal cells. This encoding strategy, and the structure of the signal itself, means variations in small chirps cannot be discriminated (Allen and Marsat, 2018; Marsat et al., 2009). Conversely, both big and small chirps on high frequency beats produce heterogeneous responses and chirp variations are accurately discriminated (Allen & Marsat, 2018; Marsat & Maler, 2010).

The mechanisms of chirp production are similar in *A. albifrons* and *A. leptorhynchus*, but chirp structure differs in a number of ways. Chirp duration is the most notable difference in these species. *A. leptorhynchus* chirps are typically tens of milliseconds long, whereas *A. albifrons* chirps are generally over 100 milliseconds long (Dunlap et al., 1998; Turner et al., 2007; Zupanc and Maler, 1993). This lengthening is thought to be a recent evolutionary change, with shorter chirps representing the basal state of this branch (Smith et al., 2016). *A. albifrons* show differences in frequency tuning from *A. leptorhynchus* (Martinez et al., 2016) that may be adaptations for coding these long chirps. Additionally, *A. albifrons* chirps typically do not fall into discreet “small” and “big” chirp categories like those of *A. leptorhynchus* (Turner et al., 2007), and chirps of varying durations and frequency are used in all contexts (Kolodziejski et al., 2007). Therefore, it is unknown whether varying chirps have different neural and behavioral impacts in A. *albifrons*.

In this study, we examine coding of conspecific social signals and their behavioral use to understand if the sensory system is adapted to the specific characteristics of the communication system of *A. albifrons*, and if so, to what extent. We compare our findings to the behavior and physiology of the closely related *A. leptorhynchus* to identify specific neurophysiological adaptations that reflect differences in the structure and behavioral use of chirps in these two species.

## METHODS

### Animals

All animals were housed according to WVU IACUC standards, protocol 151200009.2. Wild-caught *Apteronotus albifrons* and *Apteronotus leptorhynchus* were obtained from commercial fish suppliers and housed in small groups (1-10 fish per tank). Tank conductivity was maintained at 200-500 μS. Sex was not confirmed, but adult animals with a wide range of EOD frequencies (600-1300Hz) were used for all experiments, indicating that we likely had multiple animals of both sexes.

### Electrophysiology

Surgical techniques were identical to those previously described in (Marsat et al., 2009; Marsat and Maler, 2010). Cells of the LS of the ELL were targeted. *A. albifrons* brain anatomy is very similar to that of *A. leptorhynchus*, so major landmark blood vessels described in Maler et al., (1991) and electrode depth served as an adequate guide to locate LS pyramidal cells (see Histology). *In vivo* recordings were made via metal filled extracellular electrodes (Frank and Becker, 1964) and amplified with an A-M Systems amplifier (Model 1700). Data were recorded (Axon Digidata 1500 and Axoscope software) at a 20kHz sampling rate. ON and OFF cells were identified using known response properties, particularly responses to sinusoidal stimulation and spike triggered average waveforms calculated from responses to 0-60Hz noise (Saunders and Bastian, 1984).

### Histology

In N=9 fish, correct electrode placement was confirmed by injection of Dextran Texas Red dye (Thermo Fisher, catalogue # D1829) at recording site using double barreled electrodes, similar to methods described in Krahe et al., (2008). After recording, dye was pressure injected with a PicoPump (WPI, PV820). Animals were then anaesthetized and respirated with a solution of Tricaine-S (.5 g/L, Western Chemicals) and perfused with 4% paraformaldehyde (Electron Microscopy Supply, # 15712) in 1X PBS. Brains were postfixed overnight, sectioned (150 μm), and counterstained with Syto59 nuclear stain (Thermo Fisher, # S11341), which allowed for clear distinction between ELL segments. In all marked sections, correct placement within the LS was observed.

### Stimuli

All stimuli were created in MatLab (Mathworks, Inc.) and sampled at 20 kHz. Stimulation was provided by a direct modulation of the amplitude (AM) of a carrier artificial EOD phase locked to the fish’s own rather than by mimicking a second EOD. This method (described below) is commonly used in similar experiments (Benda et al., 2005; Krahe et al., 2008; Marsat et al., 2009) and allows precise control over the stimulus AM. Baseline EOD was recorded via electrodes near the head and tail of the fish. Each EOD cycle triggered a sine wave generator (Rigol DG1022A) to generate one cycle of a sine wave matched to the animal’s own. This signal was then multiplied using a custom-built signal multiplier (courtesy of the Fortune Laboratory, New Jersey Institute of Technology) by the AM stimulus to create the desired modulation of the electric field. Stimuli were played through a stimulus isolator (A-M Systems, Model 2200) into the experimental tank via either two 30.5 cm carbon electrodes arranged parallel to the fish’s longitudinal axis (global stimulation) or two silver chloridized point electrodes 1 cm apart from each other positioned near the receptive field on the fish’s skin (local stimulation). The stimulus strength was adjusted to provide ∼20% contrast (the difference between the maximum and minimum of amplitude modulation divided by the baseline EOD). See Fig 1-3 for further comments on stimulus strength.

Random amplitude modulation (RAM) stimuli consisted of 30 seconds of random noise filtered using either a low pass (0-20Hz), band pass (40-60Hz) or broadband (0-60Hz) Butterworth filter. Each stimulus was played for three repetitions in both global and local stimulation configurations. Sinusoidal amplitude modulation (SAM) stimuli were 2 second long stimuli modulated at 2, 5, 15, 30, 60, and 90Hz with 2 seconds rest between each frequency, repeated at least three times. Step stimulations were 100 ms long increases or decreases in amplitude repeated for 30 seconds.

Chirp stimuli were created by using recorded EOD samples (courtesy of Dr. Troy Smith, Electric Fish Signal Library) to create a 1000Hz template *A. albifrons* EOD shape, thereby accounting for the actual EOD shape and resulting AM beat shape. Chirps of varying durations, frequency increases, and frequency rise/fall time (Table 1) were each embedded in this template EOD by decreasing the template EOD period by the inverse of the desired frequency increase for each EOD cycle in the duration of the chirp. For each 1Hz of frequency increase, the template amplitude was decreased by .11%, based on Dunlap at al. (1998). Chirps were added at the rate of 1 chirp per second. To this chirper EOD, a second EOD, either 990Hz or 900Hz, was added to create a combined EOD signal with a beat AM frequency of 10 or 100Hz. The AM of this stimulus was extracted by rectifying and low-pass filtering the combined EOD signal. The AM signal was delivered during the experiments as described above. Each chirp stimulus was played for at least 30 seconds and up to 60 seconds. Due to time constraints, chirps were only played in the global configuration.

**Table 1:** Frequency and duration properties for all chirps used. “Other” indicates differences in chirp shape not due to peak frequency or duration, such as shape of frequency rise and fall, and phase of the beat on which the chirp occurred.

| Chirp ID | Frequency Rise (Hz) | Duration (ms) | Other |
| --- | --- | --- | --- |
| 1 | 200 | 100 | $\alpha$ shape |
| 2 | 200 | 200 | $\alpha$ shape |
| 3 | 350 | 100 | $\alpha$ shape |
| 4 | 350 | 200 | $\alpha$ shape |
| 5 | 50 | 50 | $\alpha$ shape |
| 6 | 100 | 50 | $\alpha$ shape |
| 7 | 50 | 50 | Antiphase to 5 |
| 8 | 100 | 50 | Antiphase to 6 |
| 9 | 350 | 200 | Two frequency peaks |
| 10 | 350 | 200 | Ramp Shaped |
| 11 | 60 | 10 | *A. lept.* small chirp |
| 12 | 122 | 15 | *A. lept.* small chirp |

### Data Analysis

For all analyses spike trains were first binarized into a sequence of 1’s (spike) and 0’s (no spike) using a bin width of 0.5 ms. All analyses described here were performed on these binary sequences in MatLab. Statistical analyses (ANOVA, Student’s t-Test, Wilcoxon rank sum) were performed using the MatLab’s statistical analysis toolbox.

### Synchronization Coefficient

The strength of phase locking to SAM stimulation was calculated as

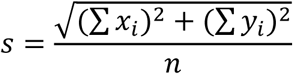

where *n* is the number of spikes in the analysis, and *x* and *y* are the cosine and sine of the stimulus’ phase at which spike *i* occurs (Goldberg and Brown, 1969; Marsat and Pollack, 2004). The vector strength, *s*, ranges from 0 to 1, with 1 being perfect precision in responding to a given phase of the cycle.

### Coherence

Lower bound (stimulus-response) coherence is a measure of linear coding of the stimulus and is calculated by comparing the spike train against the RAM stimulus (Borst and Theunissen, 1999). Stimulus-response (S and R respectively) magnitude-squared coherence (*C_SR_*) was calculated as a function of frequency (*f*) according to:

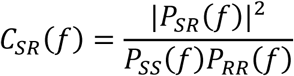

where *P* indicates the power-spectral or cross-spectral densities.

Upper bound (response-response) coherence measures the total information potentially coded in the response, including linear and non-linear information. It is measured by comparing multiple responses of one neuron to each other to determine response reliability (Borst and Theunissen, 1999). Coherence between responses *Ri* and *Rj* was calculated as:

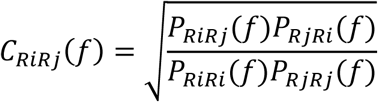

and averaged across all pairwise combinations of responses.

Envelope responses were determined by computing the lower-bound coherence between the responses and the envelope of the 40-60Hz RAM stimuli. To do so, the stimulus *S(t)* in the lower-bound analysis described above was replaced by the envelope *E(t)* calculated by rectifying and low pass filtering the 40-60Hz noise stimulus AM. Similar results were obtained when using a Hilbert transform to calculate the envelope signal.

### Burst detection

Bursts in RAM and chirp responses were identified by creating a histogram of all inter-spike intervals (ISIs) in the response. Bursting neurons have a bimodal, non-Poisson distribution of ISIs, allowing us to visually identify a threshold ISI at the upper boundary of intra-burst intervals. Spikes in groups with ISIs below the threshold were classed as occurring in bursts, while all remaining spikes were classed as tonic. This method is similar to that described in (Ávila-Åkerberg et al., 2010)

### Chirp detection and discrimination analysis

Analysis of chirps for detection and discrimination is based on Allen and Marsat (2018). Our analysis accounts for both the firing rate as well as the temporal pattern of spikes to quantify how similar or dissimilar spiking patterns are (van Rossum, 2001). For detection analysis, a window around the chirp (R_c_(t)) of length *L* (50, 105, or 205 ms) was extracted from the filtered spike train and compared to a window of beat of the same size (R_b_(t)). Different window sizes led to qualitatively similar results, and those using 205 ms are shown. The responses were convolved with an α filter, 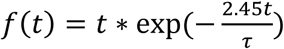, with τ being the width of the function at half maximum (3, 10, 30, and 100 ms; 30 ms shown) (Machens et al., 2003). Detection accuracy is calculated for population responses by averaging multiple spike trains:

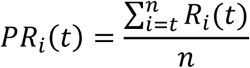

The result *(PR_i_(t))* represents a population of neurons presented with the same stimulus and mimics a neuron integrating postsynaptic potentials with similar weights. Distance *(D)* was calculated for all sets of combined responses, *PR_c_(t)* and *PR_b_(t)*, creating an array of response distances for each comparison using the function:.

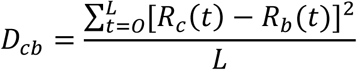

The probability distributions of the values in these arrays *(P(D_cb_)* or *P(D_bb_))* were used for analysis. Receiver operator characteristic curves were calculated by varying a threshold distance *(T)* to separate chirp and beat responses. For each threshold the probability of detection *(PD)* was calculated as the sum of *(P(D_cb_ > T))*, and the probability of false alarm *(PF)* as the sum of *(P(D_bb_ <T))*. The error level for each threshold value is *E = PF/2 +(1-PD)/2.* The values shown in Figure 3 are the minimum calculated values of *E* as increasing numbers of spike trains are included in the calculation.

**Figure 3:**
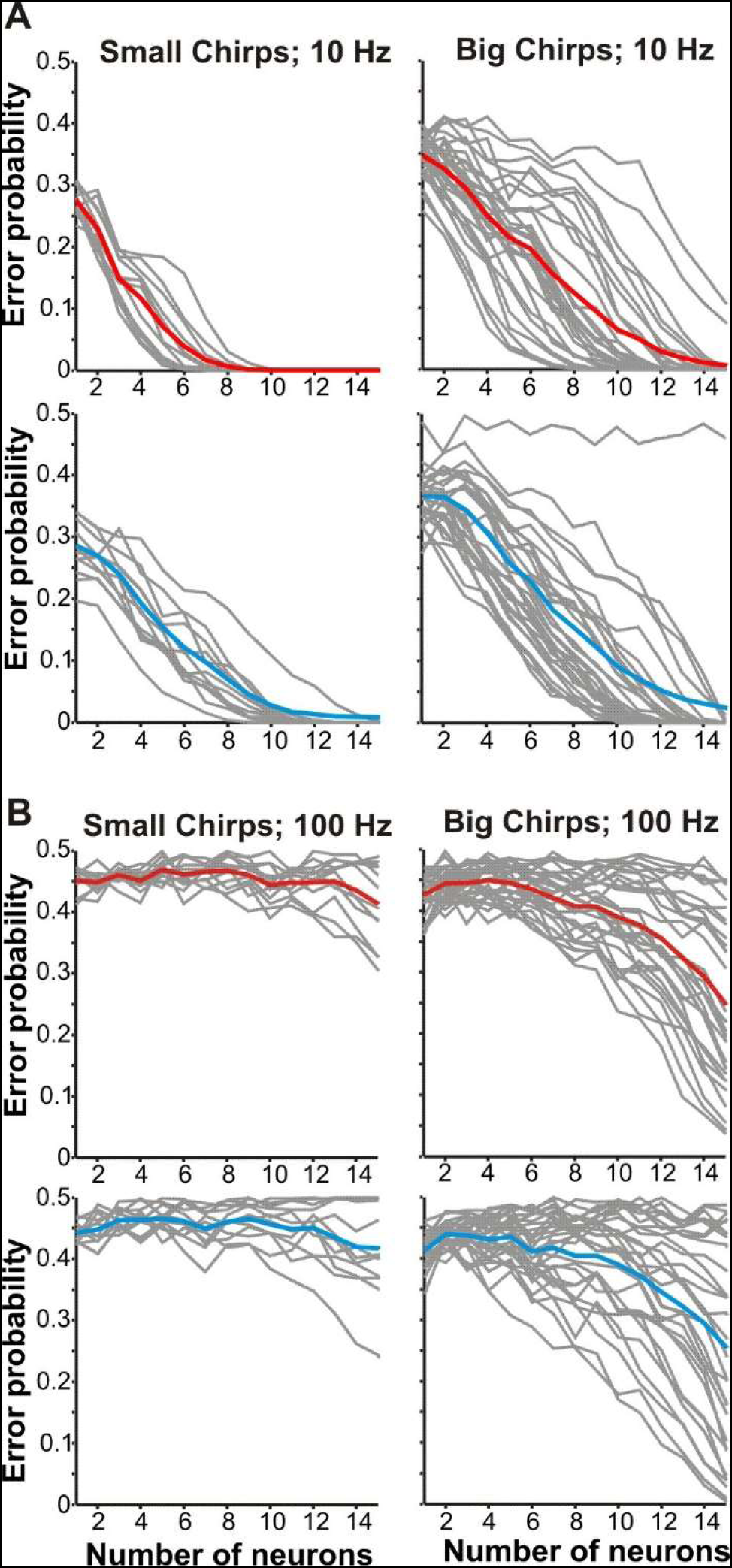
Discrimination of chirps on high frequency beats is poor. **A.** Discrimination for small chirps (Chirps 5,6,7,8, right) and big chirps (Chirps 1,2,3,4,9,10, left) on 10Hz beat. Mean ON-cell chirp discrimination shown in red, OFF-cells in cyan, and discrimination for individual chirp pairs shown in gray. **B.** Discrimination of small chirps on a 100Hz beat is relatively inefficient for both ON (red) and OFF-cells (cyan). Discrimination of big chirps varies with chirp identity, but is still poor. For chirp-by-chirp discrimination comparisons see Figure 3-1.

Discrimination analyses were similar to those for detection, but rather than comparing chirp and beat responses, the distances between two different chirps were compared. Similar to detection analysis, windows of chirp responses *(R(t))* of length *L* (50,105, and 205 ms) were used for analysis and the results using 205 ms are shown. The responses were convolved with an α filter and population averaged as described above. Up to 200 random combinations of spike trains from all recorded neurons were used for all comparisons as more become computationally prohibitive without improving results. Responses to a given chirp against a different chirp (x vs. y) were compared as well as multiple responses to the same chirp (x vs. x). Spike train metric distances (*D*_xy_ or *D*_xx_) the ensuing distribution and error rates were calculated as described for the detection analysis.

### Adaptation

Adaptation to stimuli was measured by using the MatLab Curve fitting toolbox to fit an exponential curve to a plot of instantaneous firing rate for each neuron resulting in a time constant τ. The portion of the response to use for fitting was determined by selecting the time of peak firing rate and the following 500 ms.

### Behavior

Twenty-eight behavior trials were conducted in a small tank (27 × 27 × 14 cm) containing water with conductivity, pH, and temperature matched to the home system, and one shelter tube. The tank was enclosed to block ambient light, lit with infrared lights and all trials were recorded via infrared camera (Logitech HD Pro Webcam C920). 14 cm carbon rod electrodes placed diagonally from each other in each corner of the tank recorded electrical activity, which was then amplified (A-M Systems, Model 1700) and recorded using a computer sound card.

Stranger fish from different home tanks were paired. Defining physical features (size, markings) and EODf were noted to avoid repeatedly testing the same pairing. Individual fish were used no more than three times, with at least seven days between each trial. One fish was selected and allowed to acclimate to the test tank for 20 minutes before the introduction of the second fish. Recording began immediately upon introduction of the intruder fish. Interactions were recorded for five minutes.

To detect chirps, we used a custom MatLab script to create a spectrogram of the electrical recordings to identify individuals and mark chirp times. Chirps were visually identified as >10Hz abrupt frequency increases.

## RESULTS

### Diversity in conspecific chirp responses

In multiple gymnotid species the lateral segment (LS) of the ELL serves as the primary location for encoding of communication and social signals (Marsat et al., 2009; Metzner, 1999; Metzner and Juranek, 1997), thus we targeted our recordings to pyramidal cells in that segment. To characterize the pattern of responses of *A. albifrons* pyramidal cells we played a series of chirps mimicking the natural range of reported *A. albifrons* chirps (Turner et al., 2007). Additionally, for comparison we used a small number of chirps with properties more typical of *A. leptorhynchus* (Dunlap et al., 1998; Zupanc and Maler, 1993). Detailed descriptions of chirp properties used are located in Table 1. In addition to playing chirps with diverse properties, we also varied the frequency of the beat, presenting chirps on both low (10Hz) and high (100Hz) frequency signals.

Figure 1 displays a representative selection of the diversity seen in chirp responses for both ON-type and OFF-type pyramidal cells. Qualitatively, responses appear to be heterogeneous with variability between inhibition, excitation, and desynchronization. On high frequency beats (Fig 1A) we see a transient increase in tonic firing rate in OFF-cell while ON-cells are briefly and weakly inhibited during the chirp. On low frequency beats (Fig 1B), the responses also consisted of brief increases in firing rates, but were additionally characterized by periods of inhibition lasting the entire duration of the chirp. We did not observe synchronized burst firing across the population that is typical of *A. leptorhynchus’* responses to small chirps on low frequency beats (Marsat et al., 2009; Marsat and Maler, 2010), the strategy that limits ability to discriminate these chirps (Allen and Marsat, 2018). Therefore, all chirps appear to be coded primarily through increases in firing among OFF-cells, and inhibition and desynchronization and cancelation of the beat (See Figure 1-1) of ON-cells. These results suggest that *A. albifrons* pyramidal cells uses the same coding patterns for all chirp types, but possibly exhibit differences based on beat frequency. Furthermore, a strong adaptation (see Figure 1-2) in firing rate seen between the beginning and end of the chirp response could cause chirps of different length to be poorly discriminated. Therefore, we next quantified how well the neural responses could support both detection and discrimination of chirps.

**Figure 1:**
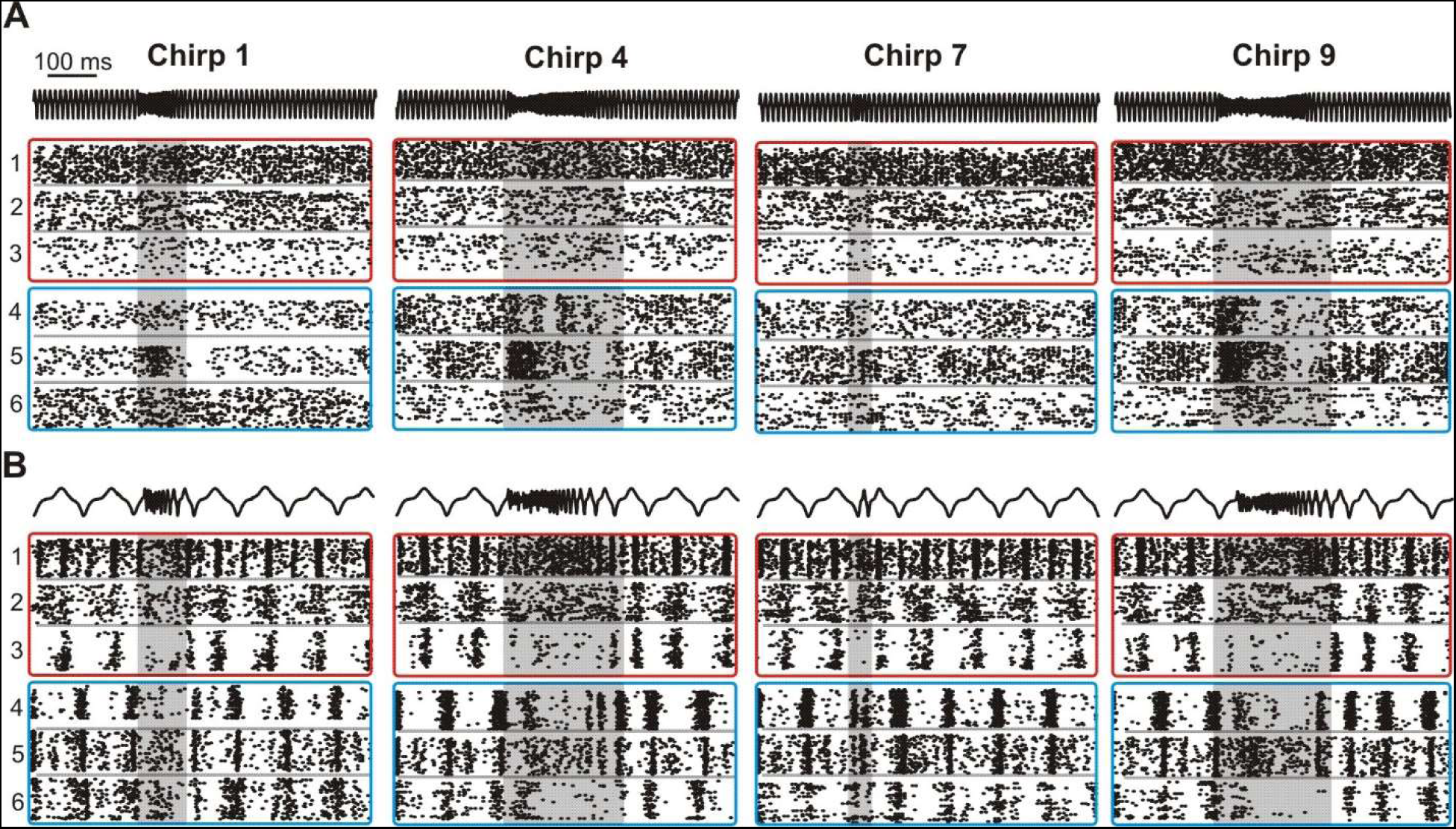
ELL Pyramidal cell responses to conspecific signals. **A.** Raster plots of chirp responses on a high frequency (100Hz) beat. AM of the EOD stimulus is shown in black; 3 representative OFF-cells are shown in cyan boxes while 3 ON-cells responses are displayed in red boxes. The same 6 neurons are used for all panels. **B.** Responses to chirps on a low frequency (10Hz) beat. For a detailed description of all chirps used, see Table 1. The shaded area highlights the duration of the chirp. See Figure 1-1 and 1-2 for comments and analysis of specific response properties (adaptation time and biphasic responses to low frequencies).

### Detection of chirps varies with beat frequency

We used a spike metric distance to measure the variability in firing patterns and how reliably an ideal observer could distinguish chirp responses from beat responses. The results are presented as an error level as a function of the number of neural inputs pooled in the analysis. More efficient encoding would lead this measure to drop faster from 0.5 (chance level) as a larger population of neurons is used in the detection task.

A small number of ON-cells (about eight) are sufficient to accurately detect chirps, comparable to *A. leptorhynchus* (Marsat and Maler, 2010) (Fig 2A). However, this level of accuracy only holds for chirps on the 10Hz beat. ON-cell performance on 100Hz is extremely poor, even with high numbers of spike trains included in the analysis (Fig 2B). OFF-cells are also able to detect chirp occurrence on low frequency beats for all chirp types (Fig 2C) but on 100Hz detecting chirps was less reliable and accurate (Fig 2D). The data suggest that with a large enough sample of neurons accurate detection is possible, but less robustly than on low frequencies. Therefore, while chirps in all contexts are potentially detected, detection sensitivity is much higher in low frequency contexts.

**Figure 2:**
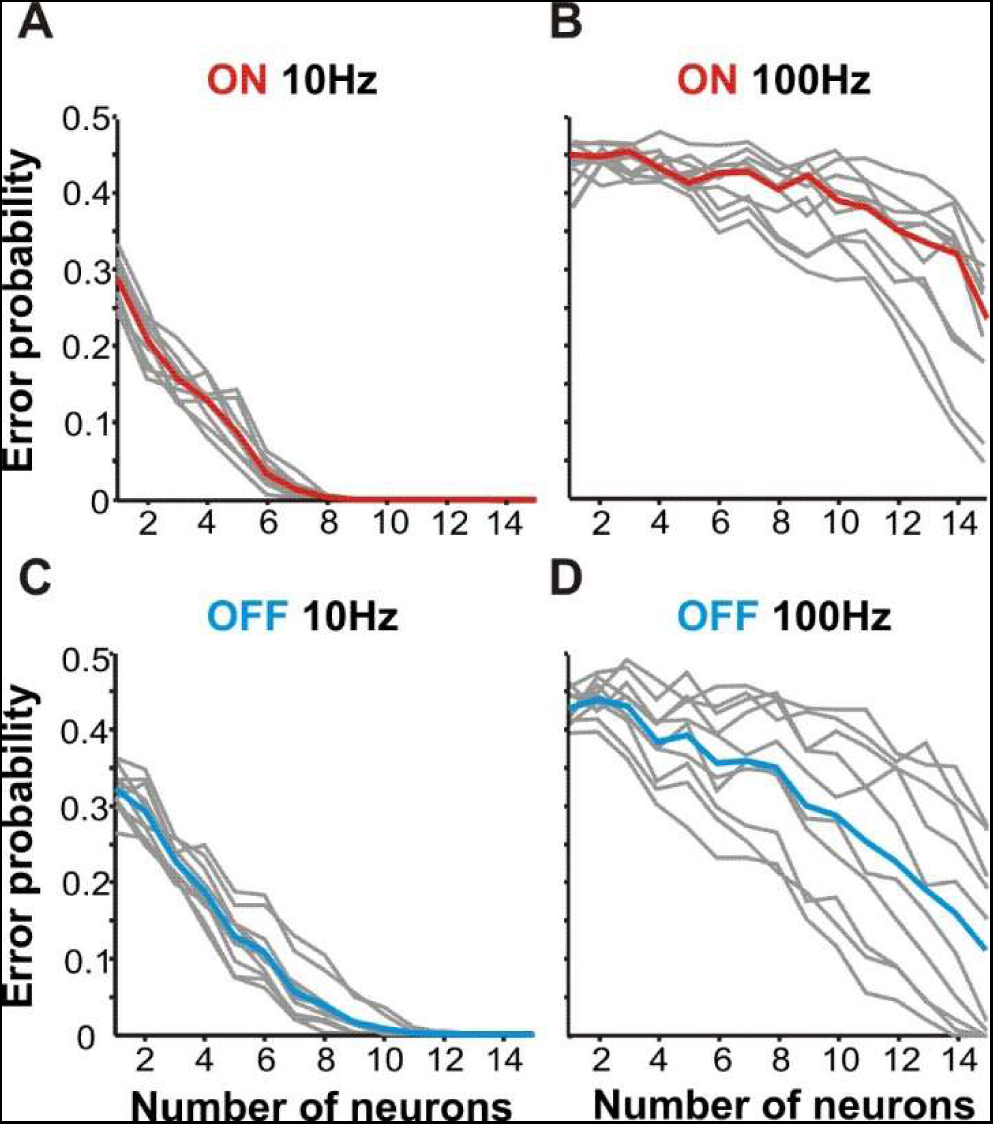
Detection efficiency depends on beat frequency. **A.** Detection of chirps on 10Hz beat by ON-cells as a factor of neurons included in the analysis (n = 17). Error probability is the probability of an ideal observer to correctly assign a spike train as a chirp or beat response. Detection error levels for individual chirp identities are shown in gray. Red line indicates mean detection error for all chirps. ON-cells can reliably detect the occurrence of all chirps. **B.** ON-cell performance is worse on 100Hz beats. **C, D** Mean OFF-cell performance (cyan, n = 16) is also more efficient on a 10Hz beat.

### Discrimination between chirps also varies

We assessed the amount of information carried by the response pattern about chirp properties that could support the discrimination of chirp variants. This analysis is similar to that used for chirp detection, but rather than comparing responses of chirps to beats, we compare responses between chirps. While *A. albifrons* chirps are not typically grouped into distinct categories, we still use the terminology used in *A. lepthorhynchus* to describe the two ends of the spectrum of chirp properties (small < 150Hz increase; big > 150Hz). Figure 3 shows discrimination ability for chirps grouped by size (See Table 1; Small chirps: 5, 6, 7, 8; Big chirps: 1, 2, 3, 4, 9, 10) for both ON and OFF-cells on high and low beat frequencies.

Chirps of all types are more easily discriminable when presented on 10Hz beats (Fig 3A) rather than on 100Hz beats (Fig 3B). Performance for chirps on high frequency beats was variable: accurate discrimination was achieved only for chirps with large differences in properties (Fig 3-1) and both ON and OFF cell responses allow qualitatively similar discrimination accuracy. These results are unexpected, considering that the same analysis in *A. leptorhynchus* shows that chirps on high frequency beats can be discriminated well using as few as six neurons, and conversely exhibit poorer discrimination on low frequencies (Marsat and Maler, 2010) where *A. albifrons* performs best. Furthermore, we do not observe an asymmetry in coding accuracy between ON and OFF cells as observed in *A. leptorhynchus*.

When looking at which chirps are well discriminated, chirp duration and the steepness of the frequency rise seem to be the most discriminable features, while total frequency increase and number of peaks are less influential on coding (Fig 3-1). However, discriminating the most easily separated chirps is prone to error and requires a larger pool of neurons on high frequency beats. These data indicate that while qualitative observations would suggest that chirp coding is similar between *A. albifrons* and *A. leptorhynchus*, there are quantitative differences in the information content about chirps within the electrosensory system. To determine which aspects of the neural responses could be responsible for changes in chirp coding, we characterized a wide variety of response properties using an assortment of stimulation protocols.

### Frequency tuning and neural coding

Encoding amplitude modulations (AM) in stimuli is crucial for the perception of social signals. As shown above, *A. albifrons* neurons respond variably to chirps, which consist of both high frequency AM signals and the low frequency changes in the contrast of the AM (i.e. the envelope). We asked whether the accuracy of detection and discrimination of chirps by *A. albifrons* ELL neurons reflects species-specific frequency tuning properties.

Communication signals cause spatially diffuse stimulation as compared to the localized electric fields of prey items. We simulate this difference by altering the geometry of our stimulating dipoles. Using a large dipole (see methods) that simulates diffuse communication signals drives both the feedforward responses from electroreceptor to pyramidal neuron and feedback pathways driven by broad receptive fields. To separate the effects of feedback from the tuning of the feedforward circuit we also stimulated only the receptive field of each cell with a small local dipole simulating localized objects or prey (Bastian, 1986; Chacron et al., 2005).

In response to simple sine waves, *A. albifrons* exhibit the highest degree of coding fidelity at low frequencies. Firing rate for each sine cycle peaks at 5Hz for all cell types and stimulation configurations (Fig 4A). Maximum phase locking also peaks at low frequencies (Fig 4B). These values closely match previously reported values (Martinez et al., 2016). Temporal coding accuracy is conveniently quantified using random noise stimuli and information theory (Borst and Theunissen, 1999). Lower-bound coherence reflects the amount of information encoded linearly whereas upper bound coherence takes in account both linear and non-linear aspects of the response (Fig 4C). Both ON and OFF cell coherence curves peak at similar frequencies in the 5-20Hz range (Wilcox Rank Sum, p = .19). However, OFF-cells are clearly low-pass, and exhibit a sharp decline in coherence to higher frequencies, while ON-cells exhibit a slightly broader lower-bound coherence (see Fig 5 and accompanying text for more details).

**Figure 4:**
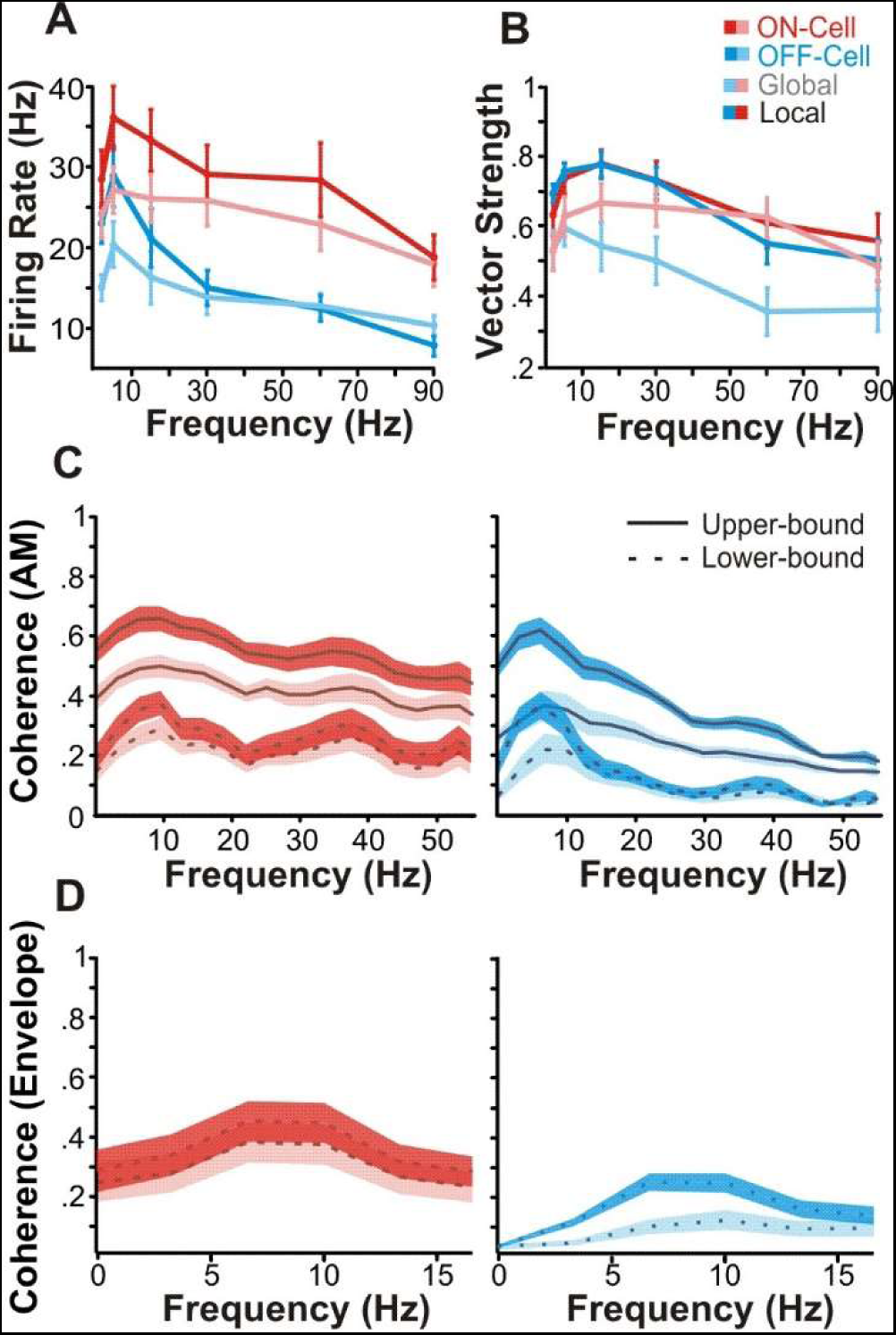
Temporal coding properties of pyramidal cells. **A.** Mean firing in response to SAM stimuli. Mean firing rate (±s.e.) for both ON (red, n = 19) and OFF (cyan, n = 15) cells in both stimulation configurations (global: pink, pale blue; local: red, cyan) peaks at 5Hz. **B.** Phase locking to AM sinusoids is also best at low frequencies. Maximum phase locking is seen at 15Hz, with the exception of locally stimulated OFF-cells, which peak at 5Hz. **C.** Mean coherence to noise stimulation is also low-pass. ON-cell coherence is shown in red, OFF-cell in cyan. The upper bound coherence measure (solid line) is based on the response-response correlations between multiple presentations of the stimulus, while the lower bound (dashed line) is based on stimulus-response correlations. Shaded areas indicate standard error; darker shading indicates local stimulation. Mean global lower bound maximums: ON-cells: 23.1Hz (±.86 s.e.); OFF-cells: 11.6Hz (±.75 s.e.), (Wilcoxon rank-sum test p = 04). **D.** Coding of low frequency envelopes is poor. Mean (± s.e.) lower bound coherence between responses and the low-frequency (0-20Hz) envelope of a bandpass RAM stimulus (40-60Hz) are displayed for both local and global stimulus configurations. Both ON and OFF cells exhibit peak envelope tuning at 10.10Hz (±1.52 s.e.) (Wilcoxon rank-sum test, p = .21). OFF-cells have noticeably lower coherence to low envelopes than ON cells.

**Figure 5:**
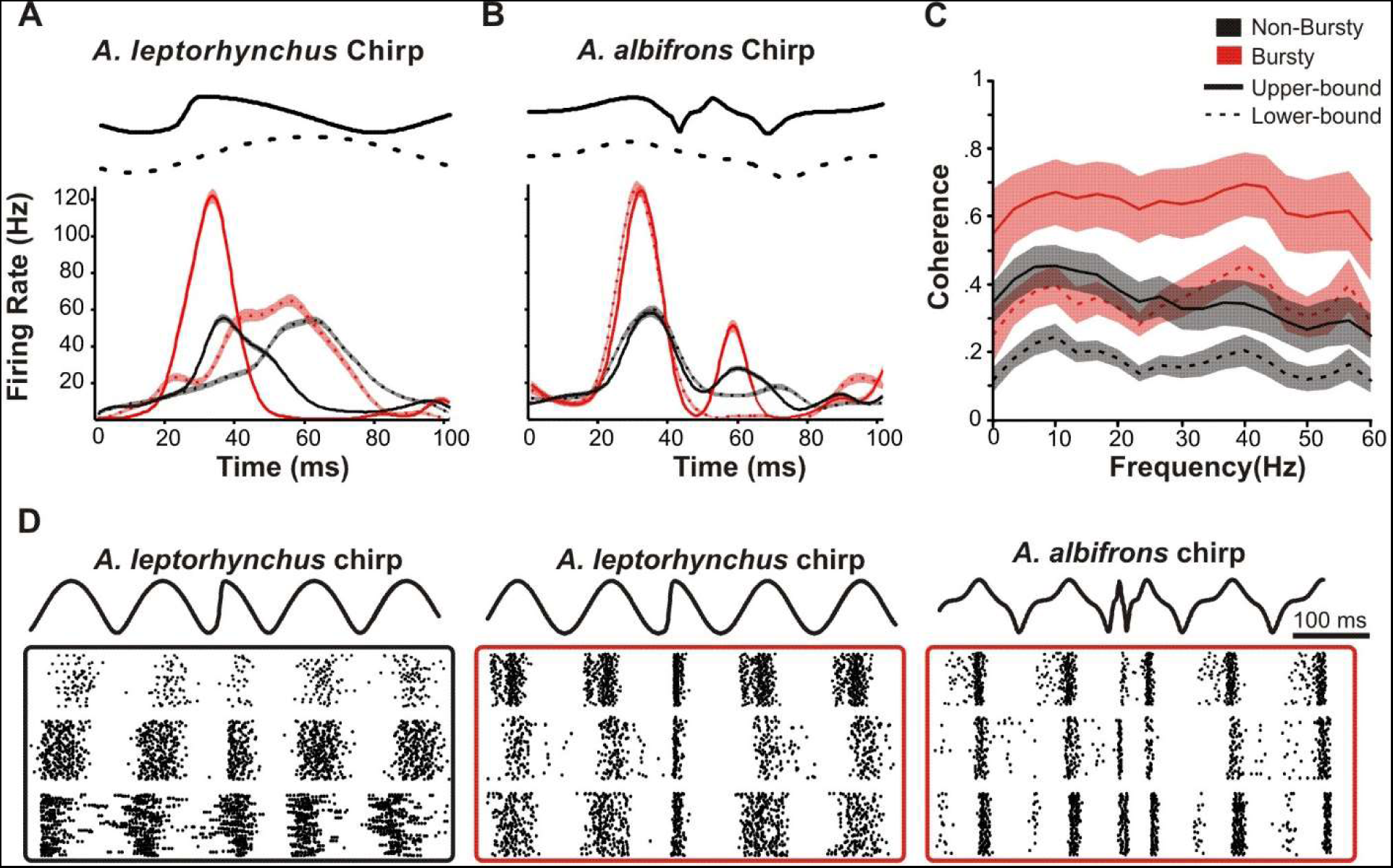
Coding of chirps by a small population of higher-pass neurons. **A.** The mean instantaneous firing rates of ON-cells over the time course of chirp and beat stimuli. The difference in peak firing rate characterize a small population (n = 5, red) that bursts in response to *A. leptorhynchus* small chirps (Beat responses, dashed lines: Peak FR 77.67Hz ± 7.11; Chirp responses, solid lines: Peak FR 159.57Hz ± 8.68; Wilcoxon ranked sum test (p < 0.001). The majority of the ON-cell population (n = 16, black) only showed a modest increase in firing (Beat Response: peak FR 62.68Hz ± 1.27; chirp response: peak FR 66.56Hz ± 4.21, Wilcoxon ranked sum test p = .002). Shaded area represents standard error. **B.** Even the bursty population fires less in response to *A. albifrons* smallest chirps compared to the beat (beat response: peak FR 155.45Hz ± 8.82; chirp response: peak FR 117.75Hz ± 9.40, Wilcoxon ranked sum test (p = .04)). **C.** Mean (±s.e.) upper-bound and lower-bound coherences for the bursting (red) and non-bursting (black) populations. **D.** Example raster plots of chirp responses used for A and B. The non-bursty population (black box) responds similarly to both *A. leptorhynchus* chirp and beat. The bursty population (red box) bursts to *A. leptorhynchus* chirps, but not *A. albifrons* chirps.). For comparable data obtained from *A. leptorhynchus* see Figure 5-1.

The movement of fish as well as the amplitude changes caused by chirps create low frequency changes in AM contrast, or envelopes (Stamper et al., 2013). During chirps, the AM will be high frequency (tens to hundreds of hertz above the beat frequency) whereas the envelope of the chirp is low frequency (e.g. 100 ms chirp will lead to a ∼ 10Hz envelope). Therefore, the coding of chirps’ low frequency content must be investigated using envelope stimuli. We characterized envelope coding using stimuli consisting of RAMs with AM frequencies between 40Hz and 60Hz, containing envelope frequencies of 0-20Hz (Middleton et al., 2006). Although pyramidal cells do encode the envelope of these stimuli (Fig 4D), the coding accuracy is low compared with the envelope coding observed in *A. leptorhynchus* (Chacron, 2006; Middleton et al., 2006; Fig 5-1). OFF-cells are particularly relevant for chirp discrimination. This is because chirps consist of decreases in envelope amplitude, which are best encoded by OFF-cells (Marsat and Maler, 2010). The observed poor coherence to envelope stimuli, particularly for OFF-cells in receiving global stimulation, may in part cause the poor chirp discrimination we observe.

### Burst Firing and Chirp Detection

As described in Marsat et al. (2009) burst firing serves an important role in *A. leptorhynchus* for the efficient detection of extremely short chirps. As qualitatively shown above in Figure 1, no conspecific chirps reliably produced a burst response in *A. albifrons.* This implies that *A. albifrons* may not be using this dedicated code for detection of very brief chirp signals. In *A. leptorhynchus* the coding of small chirps is directly related to beat frequency. On low frequencies, small chirps are shorter than the period of one beat cycle. As beat frequency increases and the period becomes shorter, small chirps begin to last longer than one cycle. This change in duration relative to beat period mediates a change in how small chirps are coded (Walz et al., 2014). In *A. albifrons* all chirps are much longer than *A. leptorhynchus* small chirps, spanning more than one beat cycle even on low frequencies, which may eliminate the need for a feature detection code specifically for extremely short chirps. Thus, the observed lack of bursting may result from the change either in signal structure (duration relative to beat cycle) or from underlying changes to physiology and bursting capability of LS neurons.

To determine if this change in signal coding is a result of signal structure or underlying physiology, we stimulated *A. albifrons* fish with *A. leptorhynchus* chirps known to elicit a burst response in 73% of *A. leptorhynchus* LS ON-cells (Marsat et al., 2009). We determined burst threshold ISI for each neuron by plotting ISI distribution of chirp and beat responses and manually selecting a threshold that best distinguished between the two (Martinez-Conde et al., 2002) in order to maximize our measure of chirp-specific bursting. The mean burst threshold was 8.0 ms ± 0.7 ms (s.e.), comparable to the burst thresholds determined for our RAM analyses and in other reports (Ávila-Åkerberg et al., 2010; Chacron and Bastian, 2008; Marsat et al., 2009).The majority of neurons increased firing rate only slightly more in response to *A. leptorhynchus* chirps than to the beat. There was, however, a small percentage (∼ 20%) of ON-cells that did reliably burst more in response to *A. leptorhynchus* small chirps than to the beat (Fig 5A, C). Most importantly, even the bursty neurons did not respond to the shortest *A. albifrons* chirps with bursts (Fig 5B, D). Unlike *A. leptorhynchus* small chirps, which may cause either amplitude increases or decreases, depending on chirp time in relation to beat phase, *A. albifrons* chirps are all long enough to span more than one beat cycle. This chirp length in relation to the beat period means that the AM waveform caused by chirps is less variable and the response they elicit is largely independent of the phase at which chirps start. This may mean that having a chirp-invariant coding mechanism, such as bursting, is not necessary in this species.

The low proportion of neurons that burst in response to *A. leptorhynchus* chirps demonstrates that there are physiologic differences between these two species in regards to burst coding of communication signals. We examined the differences between bursty and non-bursty neurons in more detail by analyzing separately their response to RAM stimuli. The subset of bursty neurons have broader AM tuning (Fig 5C) as tested by comparing the ratio of coherences summed over the 0-30Hz vs 30-60Hz (bursty cells mean ratio ± s.d.: 0.91 ± 0.12; non-bursty cells mean ratio: 2.35 ± 1.72; t-test not assuming equal variance, p = 0.0047; similar results were found using the upper-bound coherence). While most ON-cells exhibited peak coherence at frequencies between 10-20Hz (non-bursty cells mean peak frequency ± s.d.: 19.6 ± 18), these cells had significantly higher peak frequencies as high as 40-50Hz (mean ± s.d.: 40 ± 13.9Hz; t-test, p = 0.023) and exhibited high coherence across the range of stimulus frequencies. The remainder of the ON cells displayed the low-pass tuning described in Martinez et al. (2016). These data show that while a subset of ELL neurons are capable of bursting in response to short chirp stimuli, they make up a small percentage of the overall ELL population and bursting does not seem to play a role in conspecific chirp coding.

### Feature Detection

Burst firing may not be a significant aspect of communication coding in *A. albifrons*, but other uses for burst coding may be conserved. During spontaneous activity, we observed a baseline firing rate of 13.54Hz (±1.37 s.e.) with 17.01% (±2.55 s.e.) of spikes in bursts. Thus, the pyramidal neurons of *A. albifrons* are physiologically capable of bursting. In *A. leptorhynchus* burst firing can reliably signal the presence of spatially localized prey-like stimuli (Gabbiani et al., 1996; Oswald et al., 2004). We examined bursting in response to local RAM stimulation (Fig 6A) to determine if bursting could serve similar prey detection functions in *A. albifrons*. The ISI histogram of the responses clearly showed that the neurons burst to these stimuli (Fig 6B). The proportion of spikes occurring in bursts was as high as 63.07% (±5.64 s.e.) for stimuli in the local configuration (Fig 6C). The average stimulus waveform triggering burst vs single spikes follows the pattern observed in other species: slower AM for bursts than for single spikes (Fig 6D). However, in response to RAM stimuli, unlike *A. leptorhynchus* there was not a large difference in coding error between bursting and tonic spiking (ANOVA, p = .10) (Fig 6E). These data suggest that while bursts encode low frequency stimuli in the LS of *A. albifrons*, they do not appear to implement a burst-based feature detection code for either prey-like stimuli or chirps.

**Figure 6:**
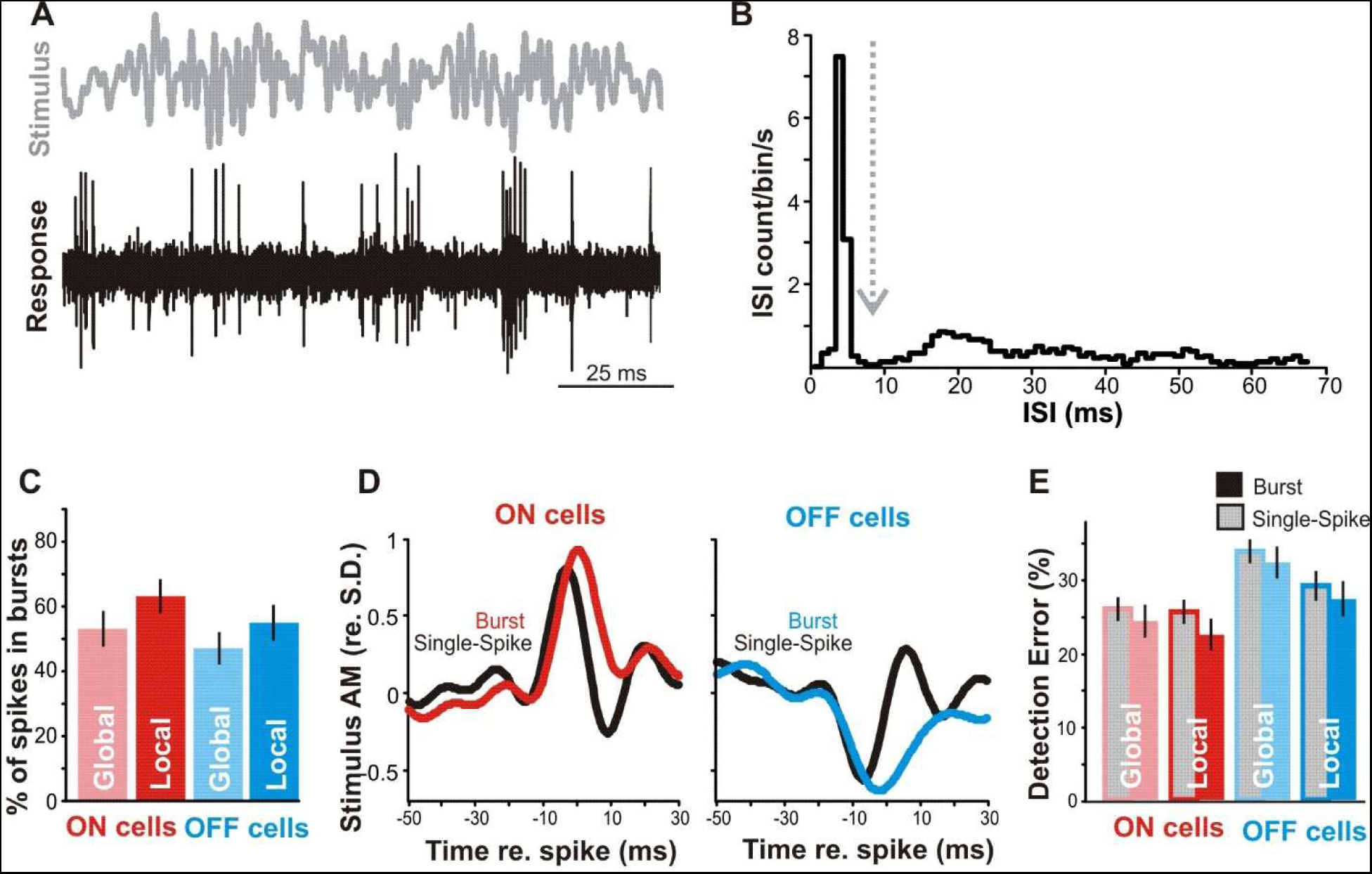
Coding of AM by bursts. **A.** Sample of noise stimulus (0-60Hz, gray) and representative spike train (black) from an OFF-cell during local stimulation. **B.** Example of ISI distribution used to determine burst threshold (dashed line). **C.** ON-cells (red) burst more than OFF-cells (blue) (ANOVA, p = .02), and local stimulation produced more bursting than global (ANOVA, p = .01). Error bars show standard error **D.** Mean burst triggered averages (red/blue) and single spike triggered averages (black) from ON and OFF-cells show that bursts are triggered by wider (lower-frequency) stimulus features than single spikes. **E.** Feature detection performance for burst and isolated spikes. In both ON and OFF cells bursts (blue/red-filled bars) tend to have lower error rates (percentage of events signaling false positives or false negatives) in detecting optimal stimulus features than single spikes (gray-filled bars) but this trend is not significant (ANOVA, p = .10).

### Chirp production and behavior

Our physiology data suggest that *A. albifrons* are able to detect and discriminate chirps on low frequency beats much more sensitively than on high frequency beats. To test this hypothesis, we recorded electrical behavior from pairs of freely swimming and interacting fish. Chirping behavior did occur but differed from that of *A. leptorhynchus* in a number of ways. Primarily, overall rates of chirping are dramatically lower than numbers reported from similar studies in *A. leptorhynchus* (Hupé and Lewis, 2008; Zupanc et al., 2006). Over 28 trials, we recorded a sum of 133 chirps, with a mean of 4.75 (±0.61 s.e.) chirps per 5-minute trial whereas in a similar context male *A. leptorhynchus* produced approximately of 125 chirps per 5 minute trials (Hupé and Lewis, 2008). The maximum number of chirps observed in one trial was 13. Due to the small number of observed chirps, we did not separate them into multiple categories for analysis, although chirps of varying frequency increases and duration occurred. Chirping frequency is correlated with the difference in EODf of interacting fish. Smaller differences in EODf correspond to higher numbers of chirps (Fig 7A). However, under our conditions chirping does not appear to be sexually dimorphic. Animals used in behavior experiments were not killed to determine sex, but we grouped them by EODf into high (>1100Hz) and low (<1100Hz) frequency groups which can correspond to females and males respectively in many populations of *A. albifrons* (Zakon and Dunlap, 1999). We observed no differences in chirp production between high or low frequency groups (Fig 7B). Further, chirp rate does not vary by pairing type, nor by relative EODf of individuals within each pair under our conditions (Fig 7B). These results confirm previous findings that chirp frequency in this species is not sexually dimorphic (Dunlap et al., 1998).

**Figure 7:**
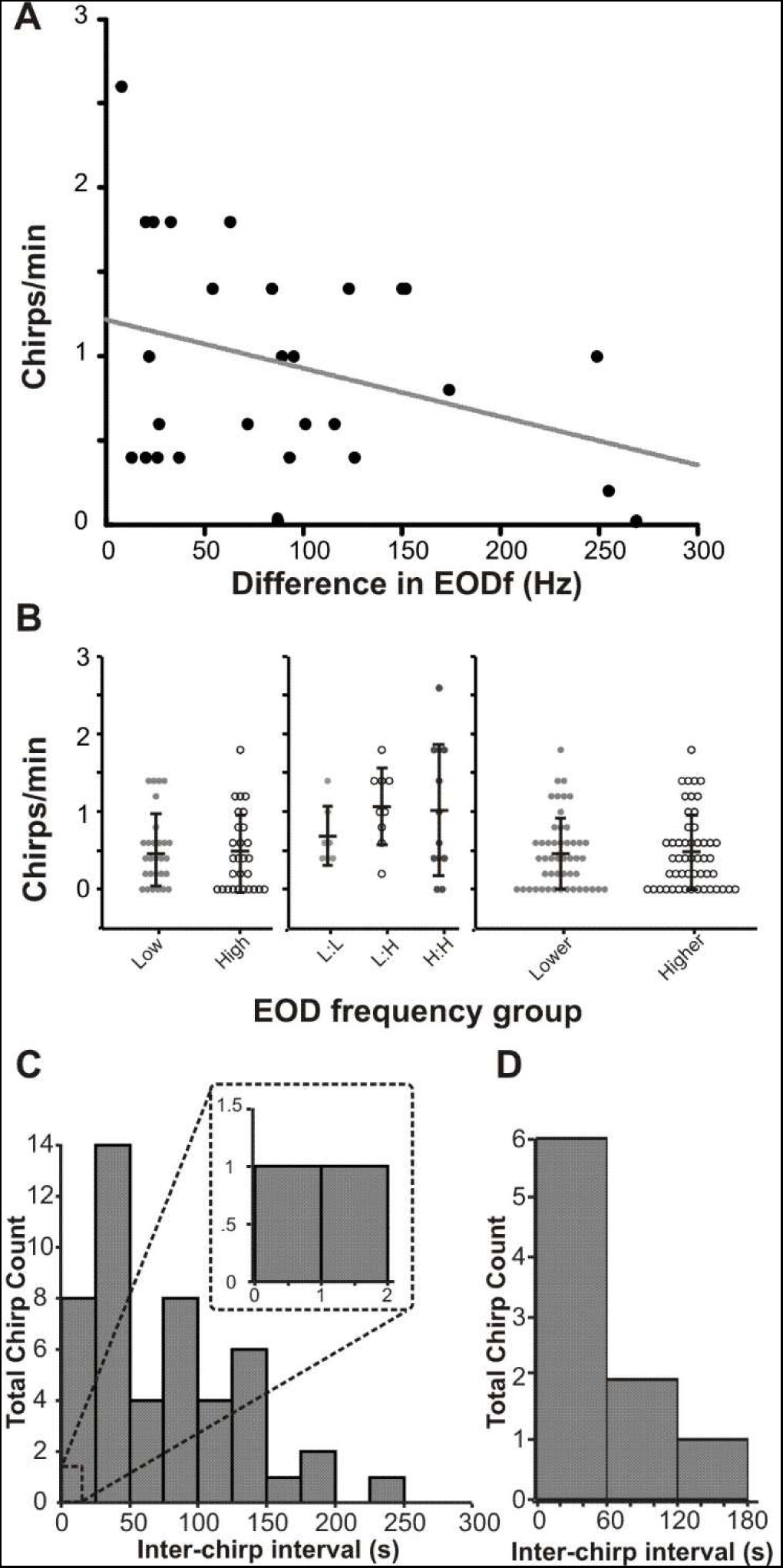
Chirping behavior in freely swimming pairs (n = 28). **A.** The number of chirps produced during interaction is correlated with difference in EODf (r^2^=.1093) **B.** Chirping does not differ by absolute EODf, EODf pair type or relative EODf. Mean chirp rate for low frequency (<1100Hz) and high frequency fish (>1100) was similar (Student’s t-test, p = .86). Mean chirps per trial based on EODf of pairing (Low: Low, Low: High, High: High) (ANOVA, p = .47), and by relative frequency of individuals within the pairing (Student’s t test, p = .55) were all extremely low and similar in all groupings. Error bars indicate standard deviation. **C.** Inter-chirp intervals between pairs binned by time fall show no echoing chirp exchanges. Inset: Enlarged section show that very few chirps occur within 2s of each other. **D.** Inter-chirp intervals distribution for individual fish follow a Poisson distribution (R^2^ = 0.9).

Chirp timing plays an important role in mediating *A. leptorhynchus* interactions as they frequently exchange chirps in an echo response pattern (Henninger et al., 2018; Hupé and Lewis, 2008; Zupanc et al., 2006), a behavior where one fish responds to the chirp of another fish by chirping back within <1s. To determine if chirp timing plays a similar role in *A. albifrons* interactions, we analyzed inter-chirp intervals for all trials in which both fish chirped. The resulting distribution of all chirp latencies between fish indicates inter-chirp latencies much longer than are typical of *A. leptorhynchus* echo responses (Fig 7C). Of all recorded chirps, only two occurred within 2.5s of each other, making an echo response unlikely to be an important feature of *A. albifrons* interactions. Furthermore, inter-chirp intervals for a given fish follow a Poisson distribution (7D) indicating that chirp production is not produced in stereotyped sequences. The lack of echoing, the long inter-chirp intervals, and the low rate of chirping in general, suggest that *A. albifrons* do not rely on the detailed decoding of individual chirp characteristics as is the case in *A. leptorhynchus*.

## DISCUSSION

We found that neurons in *A. albifrons*’ primary sensory area are able to detect chirp occurrence and do encode chirp identities on low frequency beats. However, they discriminate between chirps less accurately when chirps are presented on high frequency beats. Rather than producing discrete burst responses for small chirps and tonic responses for big chirps, all chirps in this species are encoded via graded and heterogeneous increase or decreases in firing rate. We found several adaptations in the basic response properties that may explain the neural responses to conspecific chirps. These adaptations, frequency tuning in particular, may facilitate encoding long chirps in *A. albifrons* as well as enhancing chirp coding in the behavioral context in which chirps are used most often. Therefore, we also asked whether chirping behavior could explain differences in chirp coding efficiency for high and low frequency beats. We found that the pattern of chirp use during common social interactions is very different from that of closely related species: *A. albifrons* produces very few chirps and do not respond to each other with echo chirps. Infrequent chirping is particularly pronounced in the context of high-frequency beats suggesting that certain types of interactions are not mediated by chirps. We suggest that the observed differences in chirp coding accuracy on high frequency beats may reflect a relaxation in demands for exchanging information via chirps in this context.

### Low frequency tuning could be adaptive for chirp coding

Our results confirm that pyramidal cells in the LS of *A. albifrons* are low pass. This finding parallels the low frequency envelopes produced by long *A. albifrons* chirps. However, the impact of this AM tuning on chirp coding depends on the way AMs and envelopes are processed in this system. Whereas the envelopes of chirps are low frequency, the AM are high frequency. Therefore, low frequency tuning to AM is not automatically helpful. We found that the coding of low-frequency envelopes was relatively poor, particularly in OFF cells, which-in *A. leptorhynchus*-are best at coding chirp envelopes. However, if envelope response is synthesized before the pyramidal cells (e.g. at the electroreceptors as could be the case in *A. leptorhynchus* (Savard et al., 2011)) low frequency tuning to AM could benefit chirp coding since the input to the pyramidal cells would contain low-frequencies (Metzen et al., 2018). Nevertheless, we show that the neural response does not accurately reflect the chirp envelope and that envelope coding of decreases in amplitude of RAM envelope stimuli is poor. Our data replicate the findings of Martinez et al. (2016) that also show poor coding of envelope stimuli, with coherence values of approximately 0.1. It is possible that without low frequency tuning, chirp and envelope coding would be even worse. Thus, low frequency tuning may compensate for other properties that hinder envelope coding (e.g. amount of rectification performed by the receptors; (Savard et al., 2011)).

### The role of burst coding

Bursts also often play an important role in coding specific features of communication signals (Creutzig et al., 2009; Fujimoto et al., 2011) as it is the case for *A. leptorhynchus* where bursting enhances detectability of small chirps on low frequency beats (Marsat et al., 2009). *A. leptorhynchus* small chirps are unusual in that, for low frequency beats, they span less than a full cycle of the beat. Consequently, the same chirp can be perceived as a sharp decrease in amplitude, a sharp increase, or a mix of the two, depending on the phase of the beat at which the chirp starts (Benda et al., 2005). It may be because of this aspect of chirp structure and its shortness, that the sensory system of *A. leptorhynchus* uses bursts to enhance detectability rather that optimizing the coding of these chirps’ properties. *A. albifrons* is not subject to the same constraints. The perceived AM of a chirp is largely independent of beat phase since chirps span several cycles of the beat and their long duration might make them more conspicuous (Petzold et al., 2016). Using noise stimuli, we showed that burst coding in the LS is different in *A. albifrons.* Since bursts are not used to encode chirps, and because LS primarily focuses on processing communication signals, it is possible that these neurons do not need to use burst as a feature detection signaling mechanism as in other species (Gabbiani et al., 1996; Oswald et al., 2004).

### Chirp coding and behavior

Our chirp response analysis shows that all chirps are coded with graded increases and decreases in firing rate containing some information about chirp properties but, for high frequency beats, coding accuracy is inefficient at supporting both chirp detection and discrimination. Our behavioral data might shed some light on this apparent inefficiency at coding chirps across all social contexts. We show that, in our conditions, *A. albifrons* chirp relatively infrequently to meditate dyad interactions. Furthermore, we show that there is a tendency to chirp less when the beat frequency is higher. This is supported by previous findings showing that chirp production is more frequent in interactions involving low frequency beats (Kolodziejski et al., 2007) reiterating the relevance of chirp coding on low-frequency beats rather than high frequency interactions.

### Comparison to *A. leptorhynchus* and other electric fish

Our physiology and behavioral findings are markedly different than those observed in the closely related *A. leptorhynchus.* Primarily, our data show *A. albifrons* exhibit poor chirp coding on high beat frequencies, and efficient coding on low beat frequencies. This is directly in contrast with *A. leptorhynchus* that encode chirps on high frequency stimuli well and exhibit poorer coding of chirp identity on low frequency beats (Allen and Marsat, 2018; Marsat and Maler, 2010). Furthermore, we find that *A. albifrons* do not rely on the same coding strategy as used in *A. leptorhynchus* since feature detection through bursts is not involved in chirp coding. Our data clearly demonstrate that despite being so closely related (Smith et al., 2016) the sensory coding properties of the neurons encoding communication signals have diverged between *A. leptorhynchus* and *A. albifrons.* Chirps have been suggested to be short in the ancestors basal to the two species (Smith et al., 2016) and longer chirps a development of the *A. albifrons* branch. Therefore, it is possible that the differences in frequency tuning we observed in *A. albifrons* may be adaptive for coding these long chirps.

Our behavioral results suggest that *A. albifrons* also use chirp exchanges very differently than *A. leptorhynchus.* Unlike the complex chirp interactions observed in *A. leptorhynchus*, where chirps are omnipresent and play a central role in various type of interactions (Hagedorn and Heiligenberg, 1985; Henninger et al., 2018; Hupé, 2012) *A. albifrons* chirp infrequently and generally only in low frequency difference interactions (Kolodziejski et al., 2007). While *A. leptorhynchus* have dedicated uses for different chirp types (Henninger et al., 2018), *A. albifrons* use all types of chirps interchangeably for low beat frequency interactions and may not possess distinct chirp categories (Dunlap et al., 1998; Kolodziejski et al., 2007; Turner et al., 2007). The link between chirp properties and behavior in *A. albifrons* is also less clear than in *A. leptorhynchus* (Dunlap and Larkins-Ford, 2003). Therefore, we suggest that rather than having distinct coding strategies for different chirp types, they have adopted context specific coding to better encode chirps on low frequency beats where chirps are most likely to be produced.

Chirping behavior has been investigated across many species of wave-type gymnotiforms (Hagedorn and Heiligenberg, 1985; Petzold et al., 2018, 2016; Turner et al., 2007). Details of chirp encoding, however, have only been investigated in one other species, *Eigenmannia viscerens* (Metzner and Heiligenberg, 1991; Metzner and Viete, 1996; Stöckl et al., 2014). In this species, similarly to *A. leptorhynchus*, different chirp types are used to mediate agonistic and courtship encounters (Hagedorn and Heiligenberg, 1985). In this species, unlike the apteronotids, chirp coding appears to be unaffected by beat frequency, at least at the level of electroreceptors (Stöckl et al., 2014). However, the coding strategy observed in the ELL pyramidal cells is that the duration of the chirp is the primary feature encoded. This feature is coded via brief excitation, then inhibition of both ON and OFF-cells (dependent on sign of the DC component of the chirp) that signals the duration of the chirp (Metzner and Heiligenberg, 1991). This strategy is similar to the coding observed in *A. albifrons* in low frequency interactions, suggesting that perhaps *A. leptorhynchus* represent the outlier group, and burst detection of chirps is a specialization for their small chirps.

### Conditions affecting chirp coding

Our study did not test for the effects of neuromodulation on neural tuning and chirp coding. Neuromodulation can change cell response properties for different behavioral states (Harris-Warrick and Marder, 1991). Previous work in *A. leptorhynchus* shows that serotonin enhances pyramidal cell excitability and responsiveness to small chirps on low frequency beats (Deemyad et al., 2013). While we worked on adult animals, we did not determine sex or breeding status, both of which have large effects of the frequency and quality of chirps produced (Smith, 2013). It is likely that the effects of neuromodulation due to behavioral state could affect the reception and encoding of these chirps as well, altering sensitivity to chirps and possibly even coding accuracy in response to behavioral need. This may particularly influence the coding of chirps on high frequency beats, which we observed was surprisingly poor. This kind of interaction is more likely to occur in breeding contexts, so it is possible that animals in breeding condition could be better able to detect and discriminate these signals than the results we show here. We also did not investigate other environmental factors that could influence the sensory specializations of both species, such as microhabitats, prey capture, and general sociality, all of which could drive some of the adaptations seen.

Our analysis technique is widely used to quantify the encoding performance of sensory neurons (Itatani and Klump, 2014; Mouterde et al., 2017; Neuhofer and Ronacher, 2012), and is biologically realistic (Larson et al., 2009). Nevertheless, other analysis technics could be devised to improve the discrimination accuracy estimate. Weighting the contribution of the different neurons, or weighting them differently across time could be envisioned (Larson et al., 2010). Keeping the different neurons as separate dimensions in the analysis (Houghton and Sen, 2008) or combining ON and OFF-cell responses (Aumentado-Armstrong et al., 2015) could also improve discrimination, as could accounting for neural correlation in response variability (Hofmann and Chacron, 2017). Finally, various more complex decoding methods (e.g. multilayered neural net or PCA-type dimensionality reduction) could be used for analysis. Nevertheless, we do not expect the neural mechanism to differ widely across apteronotid species so our analysis provides a meaningful comparison. The strength of our measure is that it is conservative: it gives a clear account of the information available in ELL pyramidal cells with few assumptions about what the downstream networks use for decoding.

### Trade-offs between specialization and generalization

Classical neuroethology dictates that the mode of signal production and mode of signal reception must evolve in synchrony so that senders and receivers never lose the ability to exchange information (Bradbury and Vehrencamp, 2011). There are many examples of specialization of particular aspects of sensory systems to accomplish a highly specialized tasks (Endler, 1992). In the case of communication, sensory tuning for sender-receiver matching has been shown repeatedly. However, the converse may also be true. Over-specialization may come at the cost of reduced sensitivity to more general environmental signals. In such a case, it may be more beneficial to favor sensory generalization over specialization in animals that engage in social behaviors less often than their more gregarious relatives do. Indeed, there are several examples of peripheral sender-receiver mismatching that may be explained by gains in sensitivity to prey or predator signals to (Mason, 1991; Römer, 2016). While we see in *A. leptorhynchus* dedicated codes for communication signals with distinct meaning and in distinct contexts (Allen and Marsat, 2018; Marsat and Maler, 2010), maintaining that level of specificity for conspecific communication may be costly both metabolically (Niven and Laughlin, 2008) and in regards to detecting environmental stimuli apart from communication signals.

In *A. albifrons* we show a general match between signal characteristics, low frequency chirp envelopes, and CNS sensitivity to low frequency signals, but a lack of detailed coding that would allow for efficient discrimination of chirp identity in all contexts. Additional changes that would lead to a more accurate chirp coding could come at the expense of the ability to encode other stimuli. Therefore, for this species, limiting the resources dedicated to the coding of social signals in certain contexts (e.g. aggressive encounters with a fish with a high difference in EOD frequency) may allow the system to preserve or enhance sensitivity to other stimuli.

## Extended Data

**Figure 1-1:**
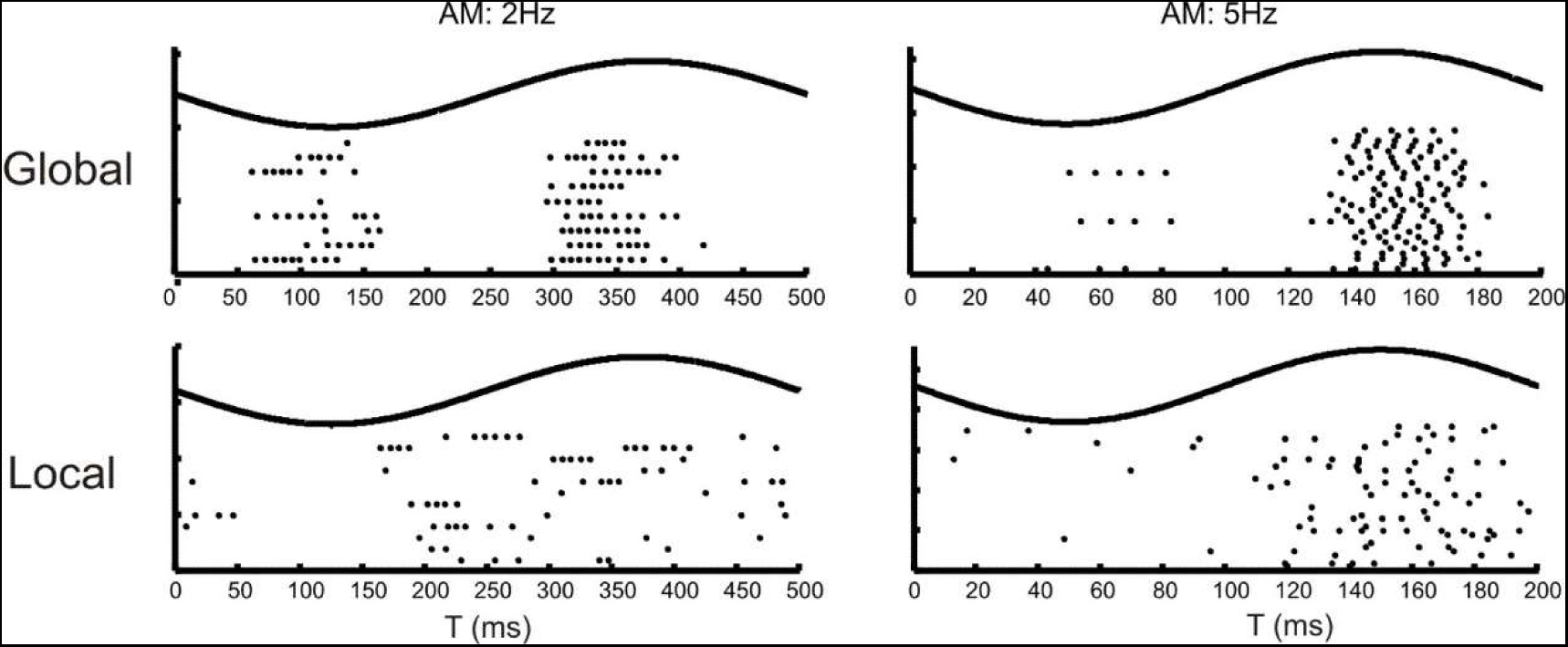
Cancellation observed in an LS ON cell. In some recordings a biphasic response to stimuli is observed. While this could indicate an overly strong stimulus, it could also indicate cancellation via feedback from the midbrain (Bastian et al., 2004). Cancellation is less prominent in the LS of *A. leptorhynchus*, but still occurs. The biphasic responses to global SAM stimuli but not to local stimulation indicate that our observed responses likely represent an increased cancellation response in the LS as a result of low frequency tuning.

**Figure 1-2:**
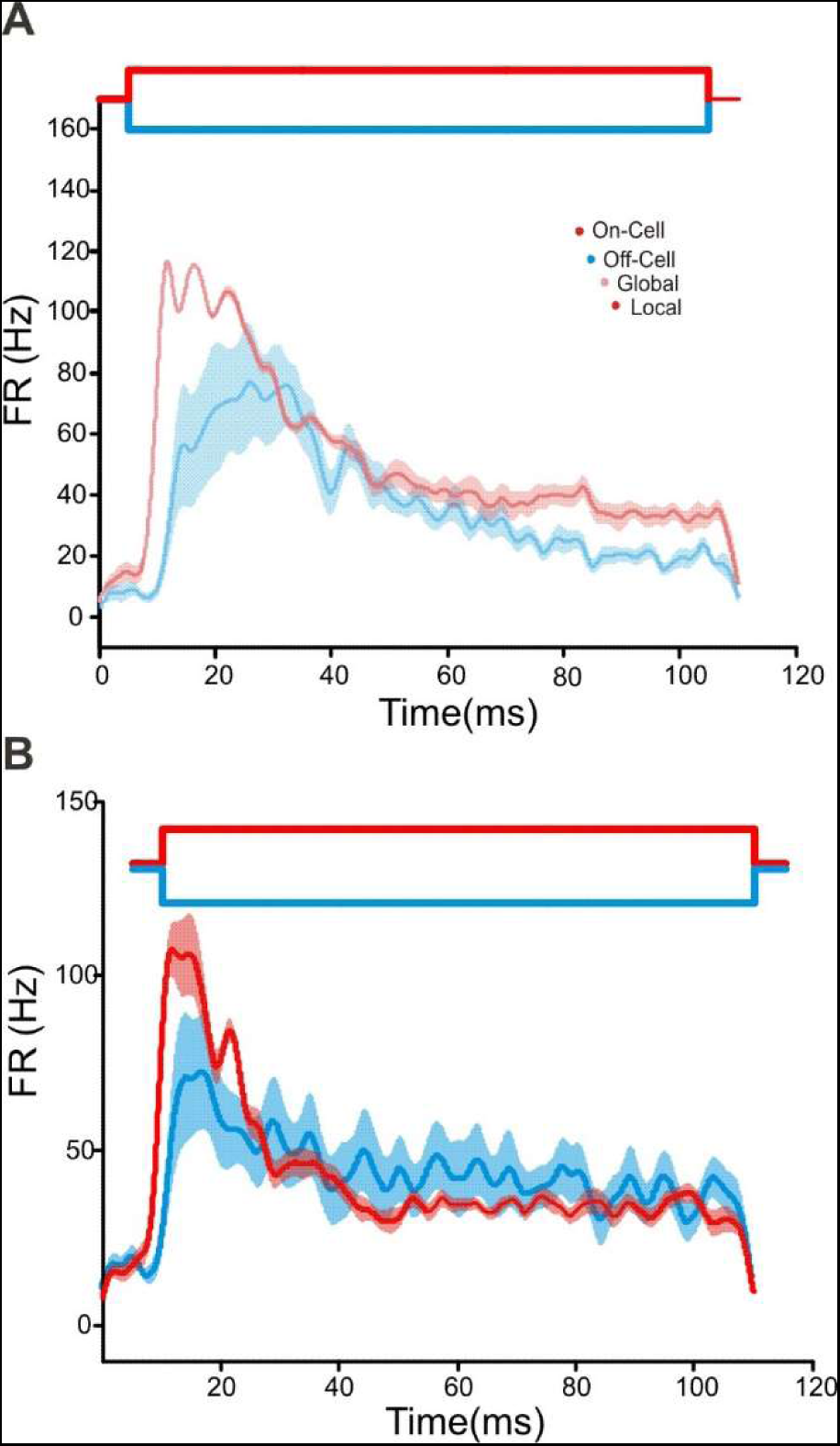
*A. albifrons* adaptation to step stimulation. A. Both ON and OFF cells’ responses decreased throughout the 100 ms stimulus without plateauing in sharp contrast with B. *A. leptorhynchus* responses to global stimuli (S2, Krahe et al., 2008 see their figure 8C&F) that adapt within 40-50 ms and then plateau.

**Figure 3-1:**
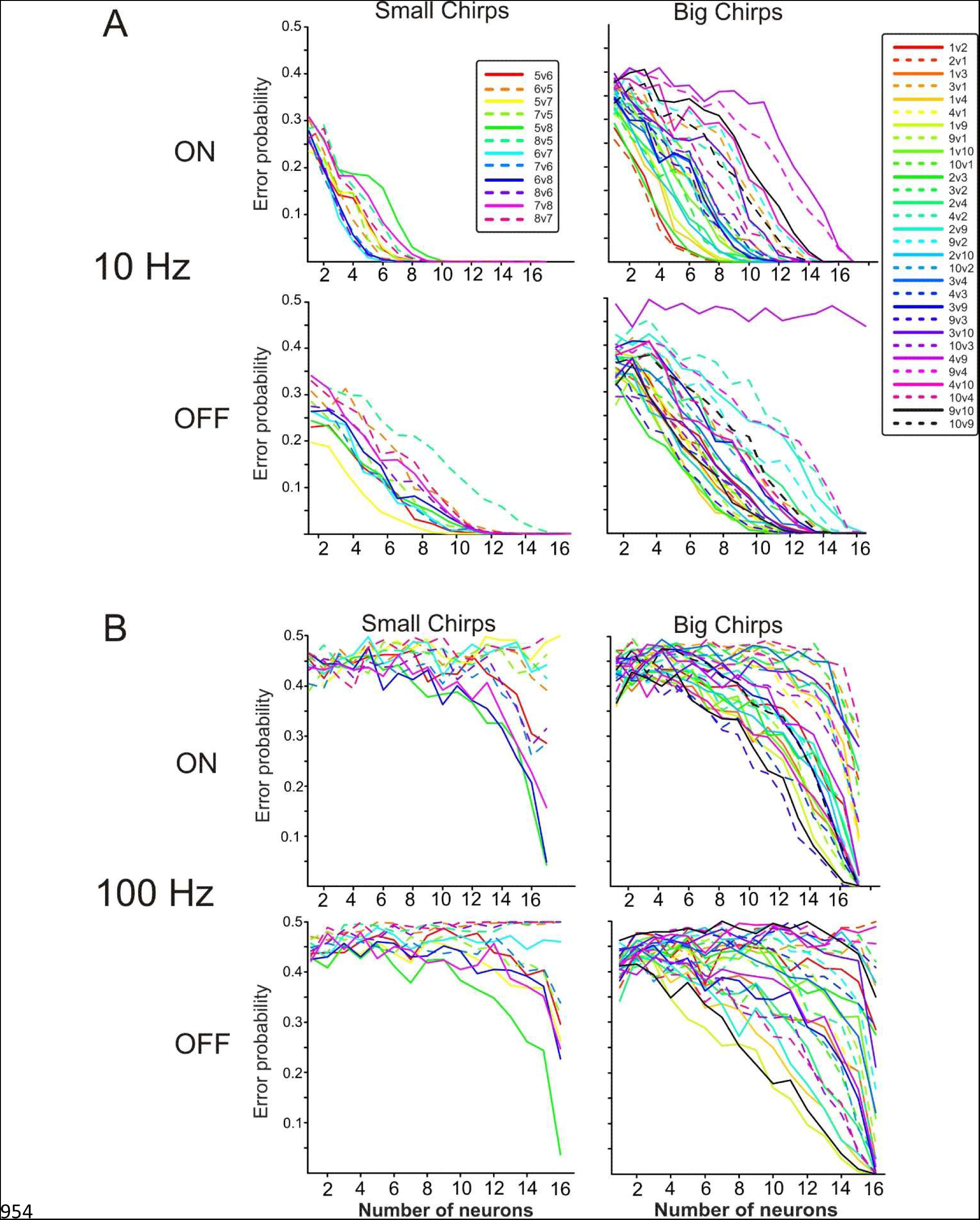
Detailed discrimination traces indicating results of testing specific chirps against each other. For ease of analysis, chirps were grouped by duration frequency and duration as either big or small. Duration appears to be the feature that is most discriminable, so big and small chirps were not directly compared. Descriptions of chirp properties are located in Table 1. A. Chirp discrimination on a 10Hz beat. B) Chirp discrimination on a 100Hz beat.

**Figure 5-1:**
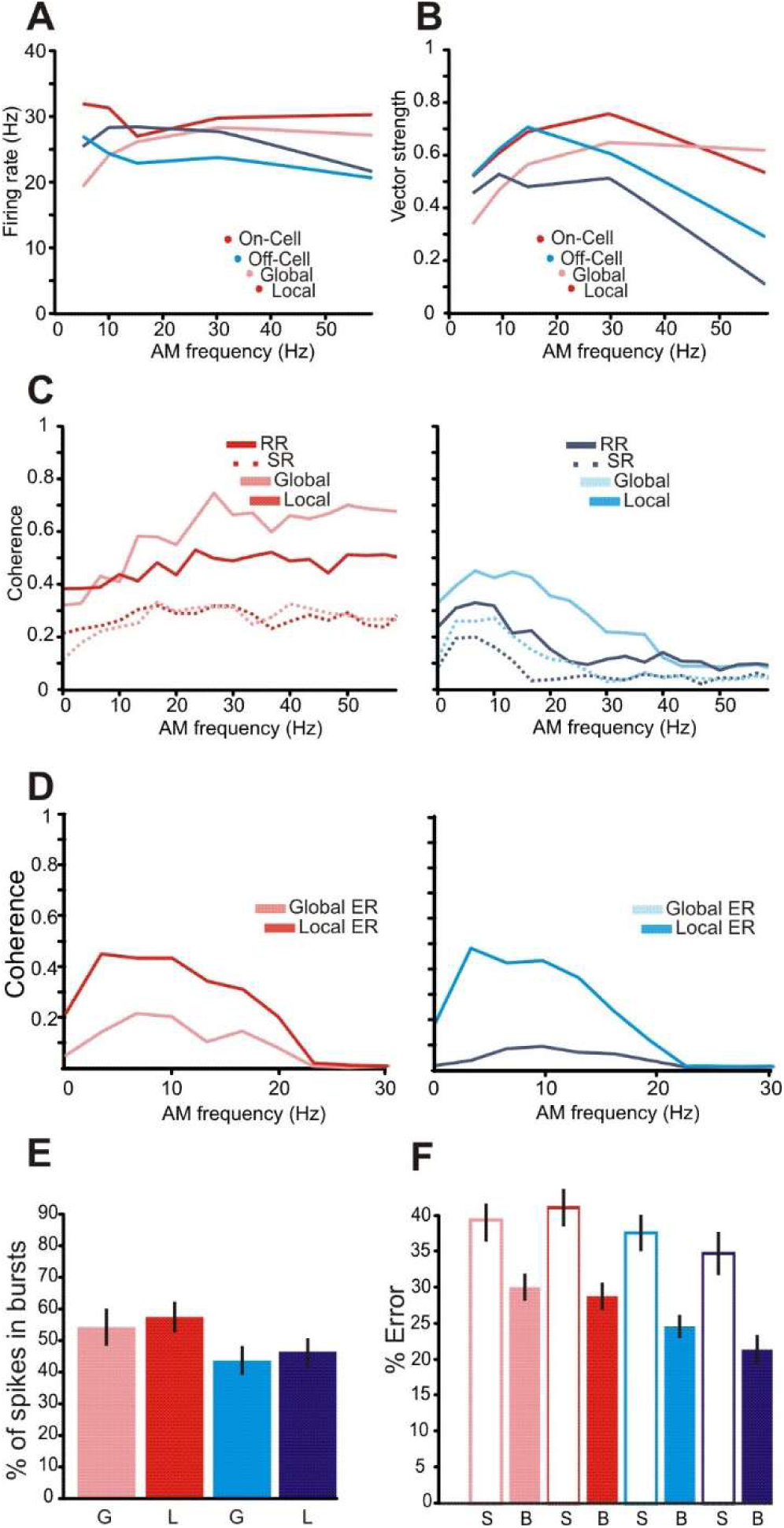
*A. leptorhynchus* data for stimulation protocols matched to those used in the main text. This data is comparable to what has already be extensively published in Krahe et al. (Krahe et al., 2008), and thus not highlighted in the main text, but indicate that our methods reproduce previously published results and indicate differences between *A. albifrons* and *A. leptorhynchus.* A. Coherence to 0-60Hz RAM stimulation. Mean coherence to noise stimulation by neuron type. ON-cell coherence is shown in red, OFF-cell in cyan. The upper bound coherence measure is shown with solid lines, lower bound in dotted lines B. Phase locking to sinusoids across 0-60Hz is best at 20–30Hz, while firing rate is relatively constant across frequencies C. Coherence to low frequency envelopes. Both ON and OFF cells perform well at coding low frequency stimulus components.

Author contributions
K.A.-Designed and performed the research, analyzed data, wrote the paper; G. M.-Designed research, created analytic tools, wrote the paper

## Acknowledgments

We thank G.T. Smith for discussions about this project, data on chirp properties and feedback on the manuscript. We thank M.J. Chacron for comments on this manuscript.

